# Sulcal pleating captures functional boundaries in human visual cortex

**DOI:** 10.64898/2026.07.20.739637

**Authors:** Anna Williams, Patricia Hoyos, Ruggaya Musa, Jesse Gomez

## Abstract

The folding of the cortical sheet within the human brain is related to its functional organization at the level of major sulci and gyri. Yet how smaller folding features hidden within the two thirds of cortex buried in sulci may relate to functional specialization of the cortex is not clear. Using human visual cortex as a test bed, we identify four hidden brain folds within the major sulcus which spans the dorsal visual stream and demonstrate that their spatial consistency across individuals is related to functional boundaries of visual cortex. In separate observation and replication participant groups, we combine structural and functional magnetic resonance imaging to show that the boundaries separating the visual field maps comprising the dorsal stream can be predicted from these hidden gyri with an accuracy matching current probabilistic definitions of visual cortex. By further demonstrating that these hidden gyri colocalize with cytoarchitectonic regions of the human brain, and that the variability in their shape and size correlates with individual differences in visual behaviors known to rely on the dorsal stream, we provide evidence for a novel neuroanatomic framework which suggests that mesoscale pleating patterns of the cortex within human sulci are functionally and behaviorally relevant.

## Introduction

Topographic organization is a unifying principle of most sensory representations in human cortex. In primary sensory regions like V1 or A1, representations follow the folding of the cortical sheet (1–3). While this structure-function relationship was thought to have only been true of primary sensory cortex, more work has demonstrated that in the ventral temporal lobe, higher level visual representations are also yoked to cortical folding consistently across brains (4–7). However, these existing models of structure-function coupling are largely limited to functional transitions that occur at macroscopic sulci or gyri, and therefore do not account for functional representations existing entirely within a single cortical fold. While models capable of predicting function from folding alone stand to advance human neuroanatomy broadly (8), no such model yet exists for retinotopic representations of visual space which populate human parietal cortex (9).

The series of retinotopic field maps emanating from primary visual cortex and extending dorsally are referred to as the dorsal visual stream (10,11). It is one of the three major visual processing streams in humans (12–14), and is involved in visual attention (9,14–16), visuomotor transformations (17–19), and visuospatial processing (20,21). It is also a frequent locus of lasting visual impairments in patients suffering strokes in parietal cortex (22–24), and likely plays a role in neurodevelopmental disorders involving reading and attention deficits. Aside from a probabilistic model built from adult data (14), there is currently no framework capable of delineating dorsal stream visual field maps in an individual brain without functional imaging and substantial expertise. There are two distinct hurdles that have likely prevented the field from defining a dorsal stream neuroanatomical model. The first is that the visual field maps comprising the dorsal stream in posterior parietal cortex exist within the paroccipital branch of the intraparietal sulcus (IPS-PO), which is a monolithic structure. Given that the visual field representations travel along a single fold, rather than traversing macroscopic cortical landmarks, current models of structure-function coupling lack the anatomical resolution necessary to account for dorsal visual stream boundaries.

Second, the field of human neuroscience has historically overlooked the nearly two-thirds of the cortical surface buried within sulci, and therefore has not accounted for cortical folding features occurring *within* major sulci. Accordingly, most atlases (25) emphasize major sulci and gyri visible from the cortical surface, yet growing evidence that smaller tertiary sulci (6,26) serve as predictive landmarks of functional boundaries underscore the functional relevance of mesoscale cortical folding. Given that intrasulcal folding features are hidden from external view, they represent a fundamental blind spot in human neuroscience. Contemporary software capable of reconstructing the cortical surface from neuroimages (27,28) now render this otherwise hidden cortex visible. The human IPS represents an ideal case study in which to determine if a major human sulcus demonstrates consistent intrasulcal folding patterns, the consistency of intrasulcal folding, and ask if these folds align with borders between functionally distinct regions (i.e., retinotopic maps). Establishing this relationship would offer a bespoke, anatomically grounded framework for identifying functional representations in populations where standard mapping is infeasible. This is especially relevant for the human dorsal stream, whose mapping relies on sustained covert attention - making it particularly inaccessible in children and clinical populations such as those with ADHD, dyslexia, or autism, for whom conventional paradigms are prohibitively difficult.

Intrasulcal folding patterns were first described in 1854 by Pierre Gratiolet (29), who noted that small folds existed within the cortex of the macaque hidden within the lunate and intraparietal sulci. He noted a consistent pleating of the cortex along the walls of the sulcus which formed small gyral ridges which he deemed *plis de passage* (PdP), likely given the appearance that they formed passages connecting the sulcal walls. While this name was translated to “annectant gyrus”, we maintain the use of their original name and refer to them as plis de passage, given that the term *plis* is closer to “pleat” which we feel better characterizes the structure of these hidden folds buried within sulci. PdP were largely forgotten until recent efforts have noted that many human sulci display this pleating phenomenon (30–32) including chimpanzees and macaques (33) but are not readily identifiable in the marmoset brain, a more evolutionarily distant primate. Recent work linking continuity of the occipitotemporal sulcus to reading (34) suggest PdP are associated with primate-specific aspects of cortical functional organization. The “hand knob” of primary motor cortex (35), a bulging of the central sulcus wall containing motor effector representation of the hand, is one example of a PdP. While the existence of PdP has been established in humans, critical questions remain unanswered. Does the human IPS display plis de passage? Are they consistent in location across brains? Do they show any relationship to functional representations of the underlying cortical sheet? Here, we combine structural MRI and population receptive field mapping through functional MRI to examine the relationship between the functional and structural topology of the human dorsal visual stream. We demonstrate four novel plis de passage within the human IPS that are not only spatially consistent across individuals, including children, but overlap with the boundaries between visual field maps defined in the same brains. Further, we show that these intrasulcal pleats predict dorsal stream field map boundaries with accuracy matching current probabilistic models. We additionally find that parietal PdP preferentially overlap with cytoarchitectonic transitions, and variability in cortical folding of this sulcal region can account for individual differences in visuospatial behavior, providing evidence for a tripartite link between structure, function, and behavior in the dorsal visual stream.

## Results

We first asked whether the intraparietal sulcus contains consistent intrasulcal folding features that could plausibly serve as anatomical landmarks for dorsal stream organization. To identify the dorsal visual stream and its relevant folds, N=33 adult participants (N=33, mean age 30.52 ± 11.71 years, N=13 female) underwent structural and functional MRI (Methods). Both T1- and T2-weighted images were collected to reconstruct the cortical surface (27,28), and all participants additionally completed three runs of retinotopic mapping for the purposes of modeling population receptive fields (pRF (36)). The pRF mapping experiment required participants to maintain fixation while attending to the sweeping bar whose contents updated at 7Hz, containing high-contrast images of ecological scenes comprised of people, objects, text, and patterns in order to drive activity in high-level neural populations (37). Attention to the sweeping bar is necessary for eliciting responses in parietal cortex (9). Functional data were resampled to each participant’s native cortical surface and each vertex’s pRF was modeled as a 2-D gaussian with an exponent fit to capture compressive spatial summation (36,38). The retinotopic maps of the dorsal visual stream were defined in each participant using polar angle reversals (9) starting at the dorsal aspect of V3 to include the following maps: V3AB, IPS-0, 1, 2, 3, 4, and 5 (**Fig 1A**). The relevant PdP traversed by retinotopic maps, as well as neighboring sulci, of the dorsal visual stream are delineated in **Figure 1B** and **Figure 2A**. For the purposes of this experiment, we will focus on visual field maps V3AB to IPS-3, which occupy the intraparietal sulcus and its parieto-occipital component (IPS-PO) just posterior to the main anterior branch of the IPS (**Fig 1B** and **Fig 2C**).

**Figure 1:**
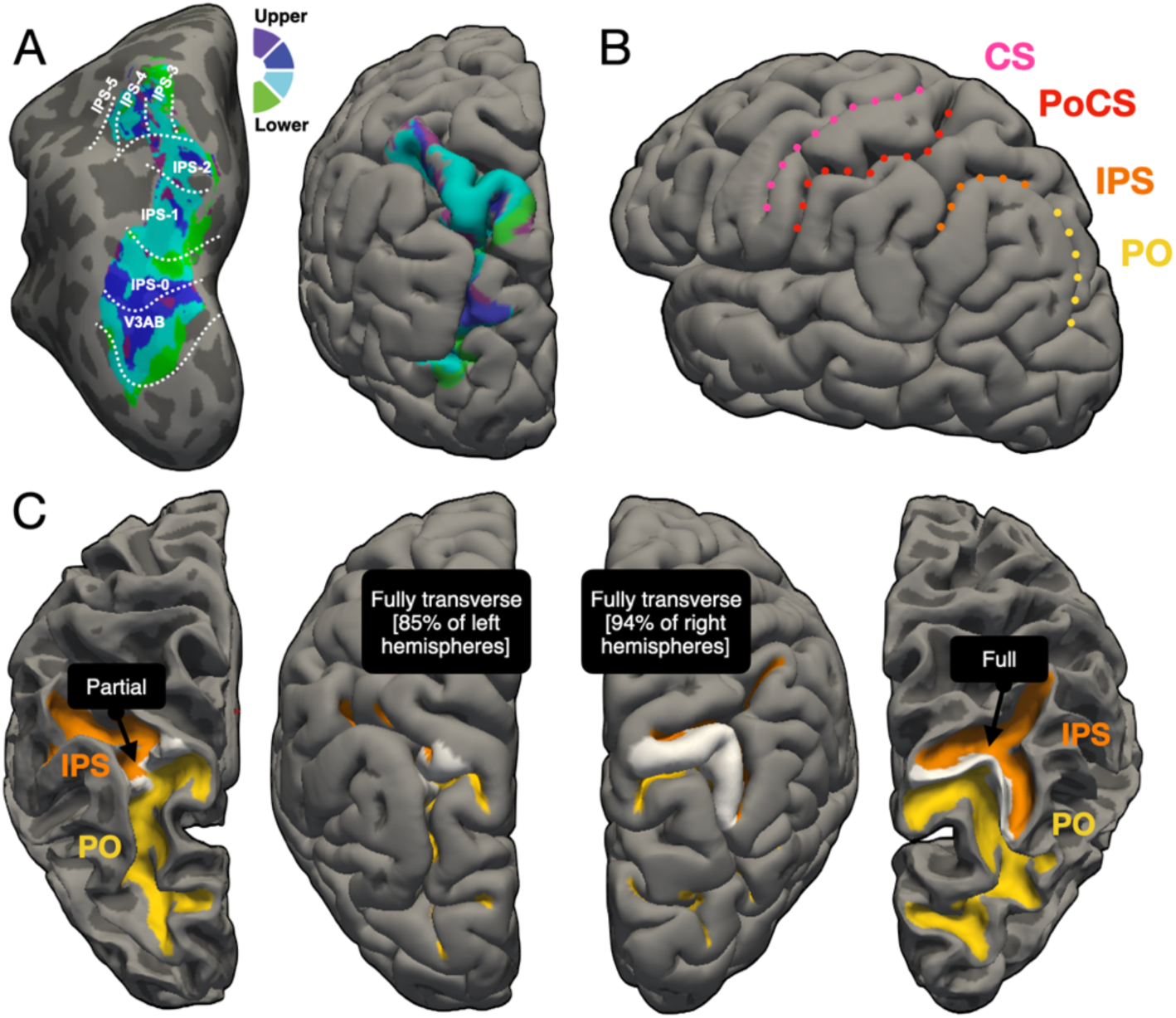
The dorsal visual stream occupies the main and parieto-occipital branch of the intraparietal sulcus. **(A)** Map depicting the polar angle position of population receptive fields at each vertex along the inflated cortical surface of a single participant. Brain on right shows majority of maps are buried within a sulcal complex. Boundaries between visual field maps delineated with white dotted lines. **(B)** Major sulci containing and surrounding the dorsal visual stream field maps. Central sulcus = CS, postcentral sulcus = PoCS, intraparietal sulcus = IPS, and the paroccipital branch of the IPS = PO. **(C)** The main and PO branches of the IPS are separated by a complete plis de passage, also known as an annectant gyrus, shown in white.

**Figure 2:**
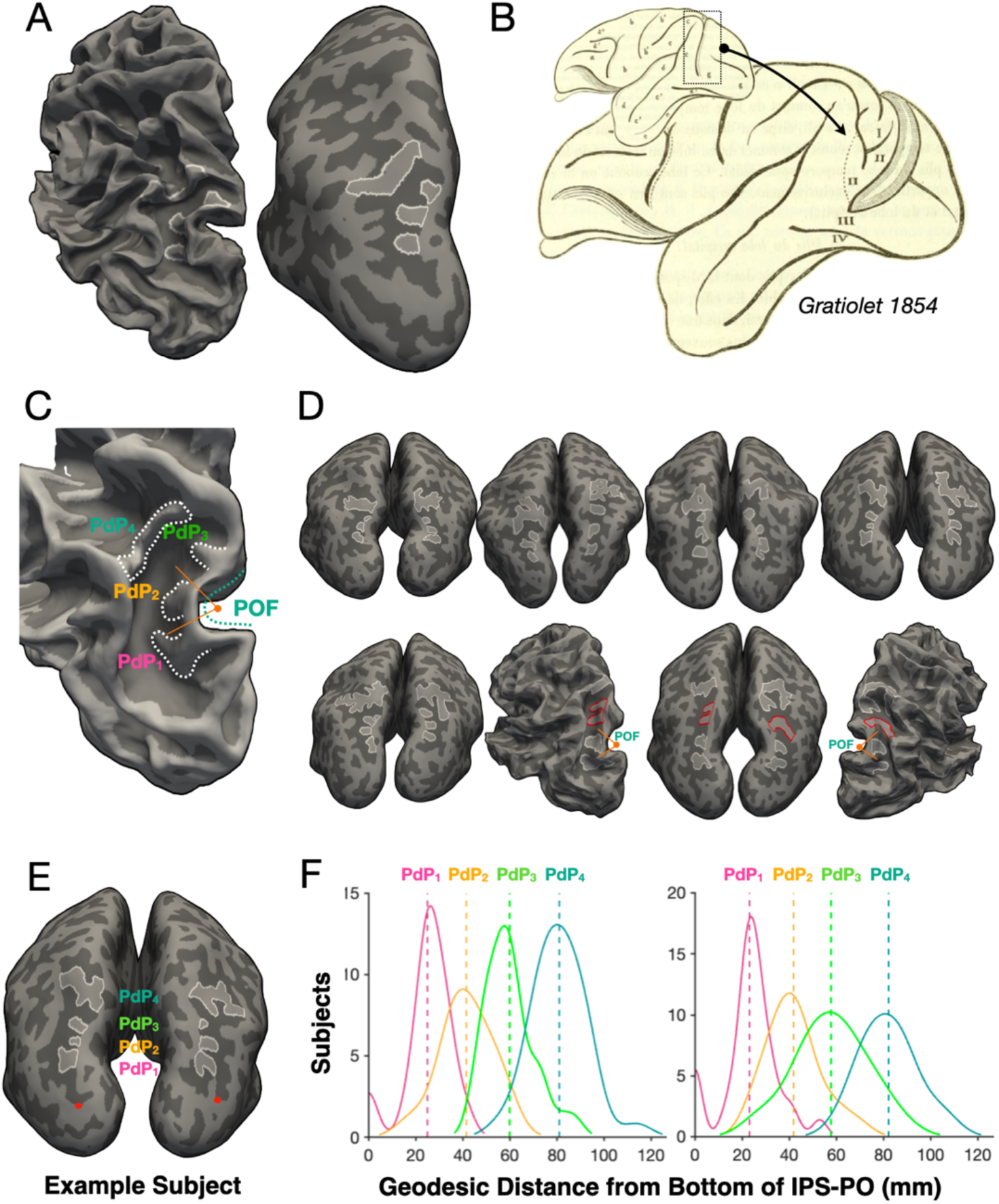
Human IPS contains three hidden and 1 complete plis de passage. **(A)** The four plis de passage of IPS-PO delineated on the white matter and inflated cortical surfaces of an example participant. The most anterior PdP is a complete gyrus that separates IPS-PO from the main branch of the IPS. **(B)** A macaque brain illustrated by neuroanatomist Pierre Gratiolet in 1854 depicts hidden PdP similarly situated between occipital and parietal cortex as in humans. **(C)** The PdP of the IPS-PO complex are numbered in ascending order from posterior to anterior. The first three separate the arc of the parieto-occipital fissure (POF) into equal thirds. **(D)** Similar to the previous panel but depicting PdP in individual participant brains. Some PdP are duplicated or completely annectant (highlighted in red), but defined and grouped consistently according to which third of the POF arc they occupy. **(E)** The posterior base of IPS-PO is marked with a red dot and used to calculate the distance of each PdP along the IPS-PO in each participant’s brain. **(F)** Histograms of the distances of each PdP to the base of IPS-PO.

Aligning with prior classification work which states that PdP can present as either superficial (visible at pial surface) or deep (hidden within sulcus) (33), we first note there are several PdP within the IPS / IPS-PO complex which vary in size and therefore visibility on the cortical surface. The continuity of the IPS from the IPS-PO was broken in a majority of hemispheres by a PdP which traversed the width of the sulcus and is visible from the surface (**Fig 1C**). This PdP interrupted sulcal continuity in 94% of right hemispheres and 85% of left hemispheres. Looking more closely in IPS-PO, we find an average of 3.13 (± 0.56) smaller and deeper PdP within the left PO and 3.13 (± 0.49) within the right PO whose depth precludes them from visibility at the pial surface (**Fig 2A**). This triplet of PdP is reminiscent of those originally described by Gratiolet in the macaque brain (**Fig 2B**), but we do not make any claims about their homology across species. In contrast to other mesoscale folds such as tertiary sulci (26), these hidden gyri were consistent to the extent that they survive the cortical surface averaging process and are visible in the FreeSurfer average brain (**Fig 3A**). The PdP of IPS-PO were consistently localized in that if one examines the IPS-PO as it wraps around the parieto-occipital fissure (POF), each pleat emerged from the medial wall of the sulcus evenly dividing the arc of the IPS-PO into thirds (**Fig 2C**). As such, we refer to the pleats within IPS-PO as PdP-1 through PdP-3. The three PdP within IPS-PO and the fourth superficial PdP (PdP-4) which separates the IPS-PO branch from IPS proper can be reliably observed in individual brains (**Fig 2D**). This regular spacing suggested that PdP might correspond to repeated functional units along the dorsal visual hierarchy. We note that occasionally a given PdP would appear as a duplicated pair, or that one of the inferior pleats (PdP-1 through 3) would fully traverse the IPS-PO, as illustrated in an example subject (**Fig 2D**). To remain consistent in our labeling, we used the arc of the POF as a guide, dividing the IPS-PO into thirds; any PdP pairs within a given third of the IPS-PO were grouped into a single pleat. Using the POF arc as a guide also aided in disambiguating PdP 1 through 3 from PdP-4 especially when multiple pleats fully traversed IPS-PO.

**Figure 3:**
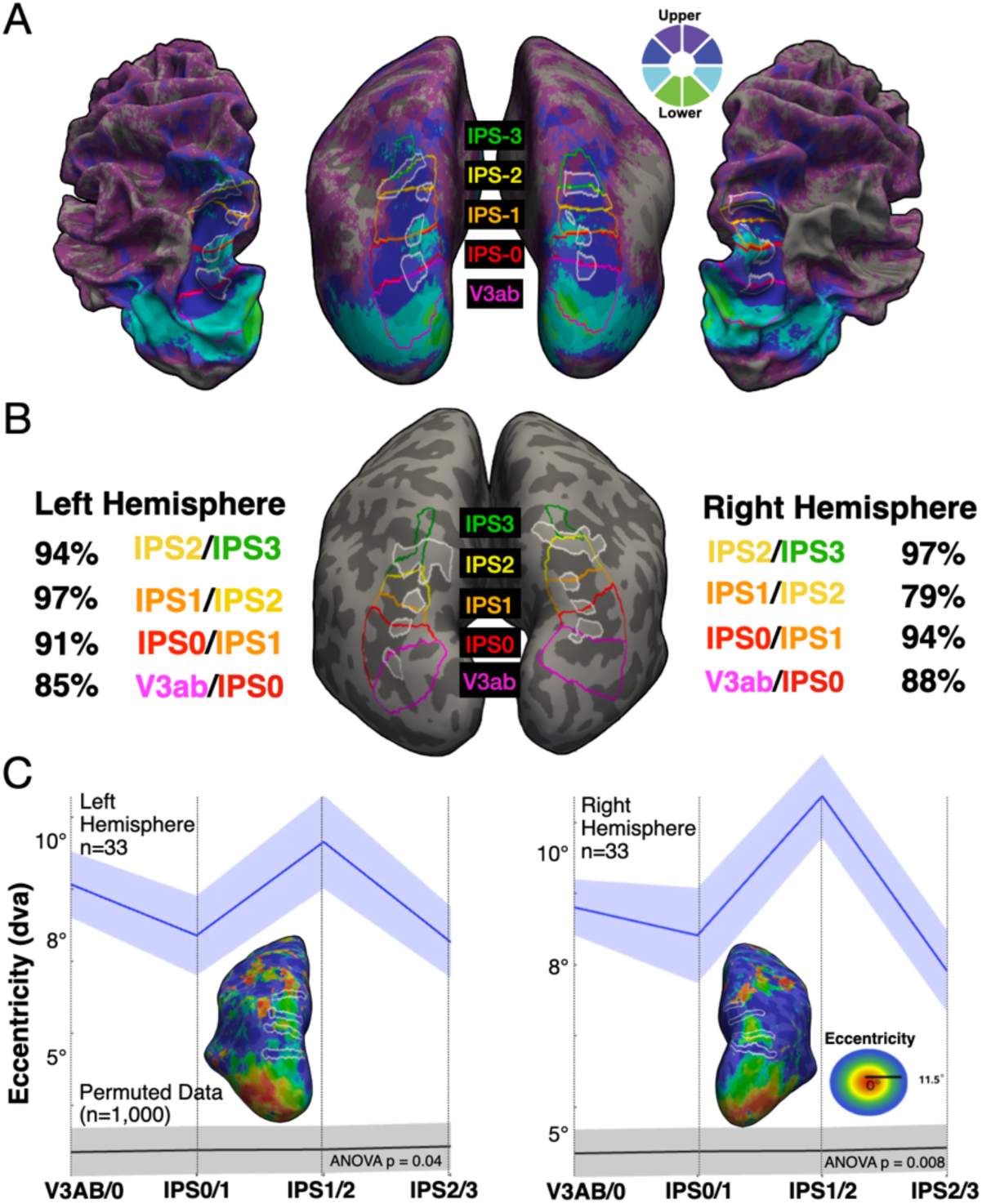
The plis de passage of the dorsal visual stream consistently align with functional boundaries associated with visual field maps. **(A)** Visual field maps defined on the FreeSurfer average brain on a polar angle map averaged across N=33 adults. The boundaries separating maps pass through distinct PdP. **(B)** Percentage of hemispheres through which a given field map boundary passes through a given PdP. **(C)** Anatomical strips shown in white defined at the location of each PdP pass through cortical regions with distinct preferences for more central or more peripheral visual space. The preferred eccentricity (distance in degrees of visual angle from center of visual space) of each cortical vertex’s pRF is shown on the inset cortical surfaces of an example participant. Mean pRF in each anatomical strip across participants is shown in blue and permuted data shown in gray.

To quantify the spatial consistency of intraparietal PdP, we measured the location of each pleat relative to an anatomical reference. On each participant’s native cortical surface, we manually identified a point located at the southernmost (posterior and inferior) portion of the IPS-PO (**Fig 2E**). We then derived the shortest path along the cortical surface interconnecting this reference point with a second manually defined point located at the center of each PdP’s gyral protrusion. From these pairs of points we quantify, in millimeters, the localization of each pleat along the intraparietal sulcus. We find that PdP are evenly spaced along the extent of the IPS-PO branch consistently across brains, almost perfectly separated by 20mm from pleat to pleat (**Fig 2F**). The distribution of PdP centers across participants are statistically different from one another (repeated measures ANOVA: Left hemisphere: F(3,31)=346.23, p < 3×10^−49^; Right hemisphere: F(3,31) = 182.87, p < 4 × 10^−38^), suggesting the locations of parietal PdP are consistent such that the variability in the location of a given pleat across brains is smaller than the separation between pleats. These distances closely matched those between PdP on the cortical surface average available in FreeSurfer (**Fig 3A**), further demonstrating their spatial generalizability in an average produced from an independent cohort of subjects. This degree of spatial regularity seems inconsistent with random sulcal pleating and may suggest a conserved folding pattern. As such, we ask if these intrasulcal pleats show any correspondence to the functional boundaries of visual representations of the dorsal visual stream.

Having defined visual field maps V3AB, IPS-0, IPS-1, IPS-2 and IPS-3 in each participant (**Fig 3A**) we can quantify structure-functional correspondence between field map boundaries and IPS-PO pleating directly. Given that prior work has observed that visual field map boundaries of the ventral visual stream occur at cortical folds (5,8,36,39), we quantified the frequency at which the boundary of two visual field maps (e.g., V3AB / IPS-0) passed through a PdP expressed as the percentage of hemispheres in which there was overlap (Boolean of map border vertices ∩ PdP vertices). We hypothesized that if PdP serve as anatomical anchors for dorsal stream organization, they should reliably coincide with visual field map boundaries. The four PdP reliably coincided with field map boundaries, overlapping in 88% of right hemispheres and 85% of left hemispheres for V3AB/IPS-0, 94% of right hemispheres and 91% of left hemispheres for IPS-0/IPS-1, 79% of right hemispheres and 97% of left hemispheres for IPS-1/IPS-2, and 97% of right hemispheres and 94% of left hemispheres for IPS-2/IPS-3 (Fig 3B). The overlap between field map boundaries is illustrated in an example brain (**Fig 3B**), as well as on FreeSurfer average cortical surface with field map boundaries drawn on a mean map of pRF polar angle fit averaged across study participants (**Fig 3A**).

While the PdP and field maps of the dorsal stream were defined by separate individuals, we next sought to offer observer-independent evidence for the correspondence of visual field representations and sulcal pleating of the IPS-PO. This analysis tests whether PdP align with the broader topography of pRF properties independent of border position. We specifically ask whether anatomical regions-of-interest defined at the center of each PdP spanning the IPS can recapitulate known changes in pRF eccentricity tuning as one ascends the dorsal stream. Retinotopic field maps of the dorsal visual stream can be grouped into pairs which share a representation of the fovea, and these pairs are separated from the next pair by cortex representing peripheral visual space. Thus, as one traverses the length of the IPS-PO from V3AB to IPS-3, cortex should alternate between a relatively peripheral bias (V3AB/IPS-0 and IPS-1/2 borders) and a foveal bias (IPS-0/1 and IPS-2/3 borders). To this end, we defined a strip of cortex including a given PdP and extending laterally to the opposing wall of the IPS-PO following the direction of orientation of each PdP in each participant’s native cortical space. Each strip included the three vertices dorsal and inferior from the center of each strip (i.e., 6 vertices in height). From this anatomically-defined ROI in each participant, we extracted the mean eccentricity value across pRF fits thresholded at 10% variance explained by the pRF model fit. We indeed observe that cortex alternates significantly between relatively peripheral to relatively more foveally-biased visual space as one travels dorsally along the IPS-PO (**Fig 3C**) in both the left (repeated measures ANOVA effect of representation (foveal vs. peripheral): F(1,32) = 4.195, p = 0.049) and right (repeated measures ANOVA effect of representation (foveal vs. peripheral): F(1,32) = 9.992, p = 0.003) hemispheres. To examine if these changes in eccentricity bias emerge simply as a property of the spacing of pleats, we permuted (n=1000) the underlying eccentricity map in each participant by rotating the map on the cortical surface to preserve spatial autocorrelation in eccentricity tuning. This spin-test control distribution does not show the same alternating eccentricity pattern (**Fig 3C**). Thus, the observed functional alternation is not explained by spatial autocorrelation alone.

Given the significant correspondence between intrasulcal pleating of the IPS-PO and functional domains of the dorsal visual stream, can pleating be used to predict retinotopic field map boundaries in a separate group of brains? This question addresses the practical utility of PdP as predictive anatomical landmarks. To this end, a separate cohort of participants (Total N=42, ages 7-20, N=30 female; Children N=21, mean age 9.24 ± 1.87, N=15 female; Adults N=21, mean age 18.71 ± 0.78 years, N=15 female) underwent structural and functional MRI similar to the prior group of participants for cortical surface reconstruction, PdP identification, and retinotopic field map delineation (40). We first note this cohort has the same number of PdP and overlap at the same frequency as the first dataset. For each participant, one researcher defined the dorsal visual field maps up to IPS-3 to serve as the metric by which anatomical predictions will be assessed for accuracy. A separate individual created roughly rectangular anatomical regions-of-interest (**Fig 4A**) to simulate visual field maps. Anatomically-defined candidate maps spanned the IPS-PO sulcus with its medial extent defined by the medial edge of each PdP, and its lateral extent defined as the vertex where the sulcal curvature of IPS-PO transitioned to gyral. The anterior-posterior aspect of each anatomically-defined map was drawn at the center of each of the four PdP along the IPS-PO. To see how these anatomical predictions of dorsal visual stream borders compare to the field’s current best model of the dorsal stream, we also aligned onto each native cortical surface a probabilistic atlas of functionally-defined dorsal stream field maps (14). This comparison directly evaluates whether anatomy-based predictions can match or exceed probabilistic functional atlases. Because the lower extent of V3AB does not yet have an anatomical definition, the probabilistic version of V3AB (8) was used in the anatomical model to define the lower boundary of V3AB. For this reason, V3AB is excluded from quantification to maintain statistical independence across datasets. We compared the performance of anatomical and probabilistic models in predicting the location of each visual field map through a signal detection framework to quantify the number of hits, misses, false alarms, in order to derive discriminability index (D-prime), dice coefficient, and Jaccard index. The field map locations predicted by the anatomical model performed significantly better than the probabilistic model across overlap measures (Jaccard, D′, and Dice) in the observation/training group from which the model was developed (repeated-measures ANOVA group effect (Anat vs. Prob): lowest *F*(1,32) = 25.109, lowest *p* = 5×10⁻^8^; **Fig 4B**). In the replication group, the anatomical model showed significantly greater Jaccard performance (*F*(1,41) = 7.660, *p* = 0.008) and numerically, but not statistically, greater performance for D′ (*F*(1,41) = 2.88, *p* = 0.097) and Dice (*F*(1,41) = 0.295, *p* = 0.590; **Fig 4C**). Another quantification of model performance, rather than overlap, is to measure the distance of each model’s field map boundaries relative to the true, functionally-defined boundaries. To this end, we calculated the vertices comprising the borders between each field map in the probabilistic and anatomical predictions. From these, we derived the centroid vertex in cortical space and calculated its distance in millimeters to the centroid of the true map border. The anatomical and probabilistic model boundary centroids were similar distances to the true probabilistic model boundary centroids (repeated measures ANOVA group effect (Anat vs. Prob): F(1,73) = 3.839, p = 0.054).

**Figure 4:**
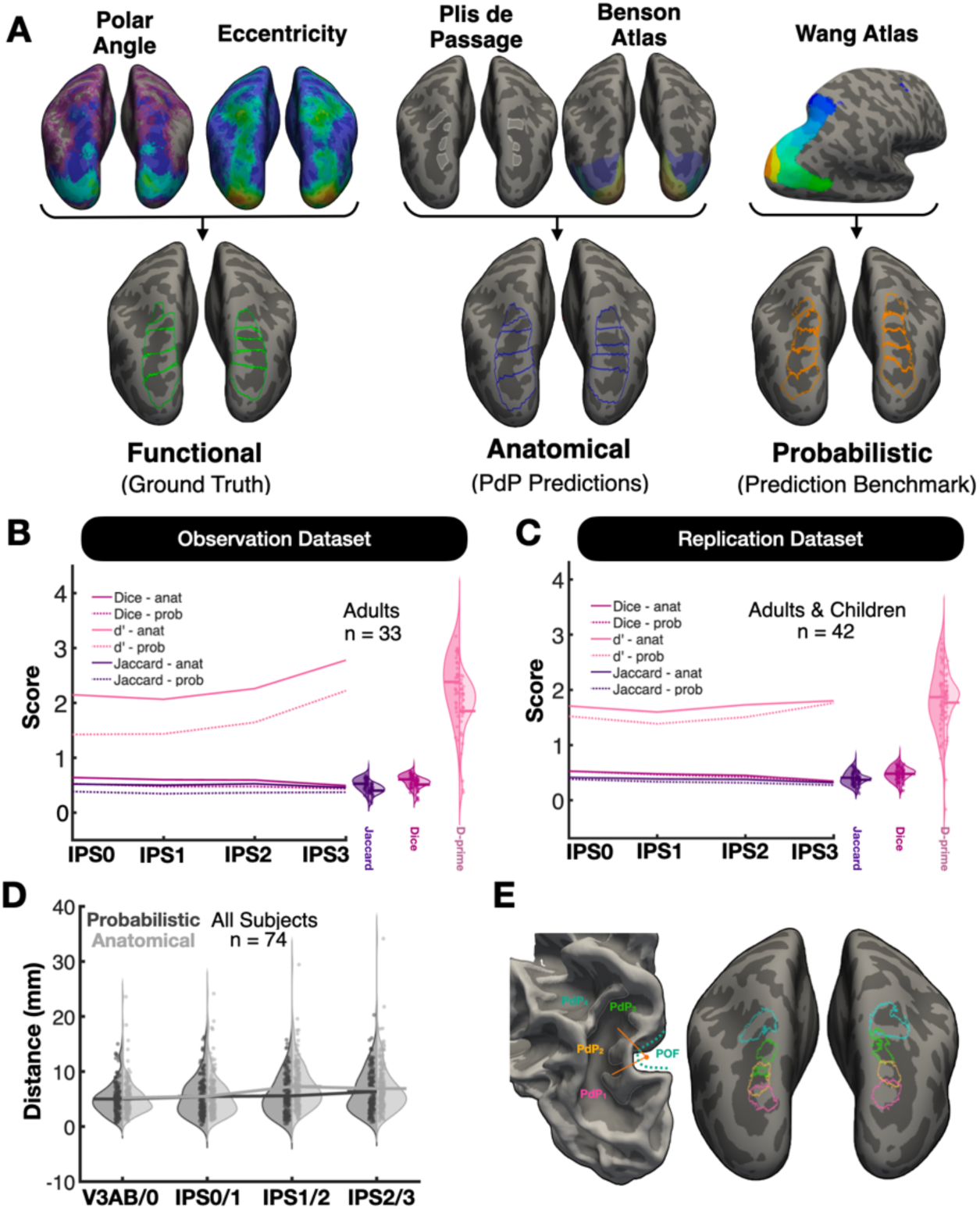
Anatomically-defined dorsal stream maps are on par with current probabilistic definitions. **(A)** Illustration for how the dorsal visual stream was defined functionally, anatomically, or probabilistically. Anatomical and probabilistic models were compared to the “ground truth” of the functionally defined maps. **(B)** Quantifications of model performance using dice, d-prime, and Jaccard index for anatomical and (darker color shades) and probabilistic (lighter color shades) definitions of field maps. **(C)** Similar to previous panel but in the separate group of N=42 participants comprising the replication dataset. **(D)** Measure of average distance of the anatomical or probabilistic boundaries separating field maps from the true functionally-defined boundary. **(E)** Probabilistic location of each PdP using binarized masks of each PdP defined in each participant’s native space and aligned to the average cortical surface.

If PdP relate to the functional differentiation of cortex, then they likely also correspond to changes in the cytoarchitecture of the underlying cortical sheet. Indeed, recent cytoarchitectonic mapping efforts in postmortem human brains show strong correspondence with visual field maps of early and ventral visual cortex (41). To address this, we again defined subject-specific anatomical strips centered at each PdP spanning the IPS-PO and ask whether these anatomical regions defined in each participant’s native cortical space show consistent and preferential overlap with cytoarchitectonic regions estimated in native cortical space from the Julich Brain Atlas of cytoarchitecture (42). The null hypothesis is that if there is no consistent relationship between PdP and cytoarchitecture, then PdP should not show any preferential overlap with cytoarchitectonic regions. We employ 7 cytoarchitectonic regions from the Julich Brain Atlas which span the IPS-PO complex (**Fig 5A**). We produced a hypothesis matrix to test the idea that PdP will show selective overlap with medial-lateral pairs of cytoarchitectonic regions (**Fig 5B**). These pairs are hIP4+hIP7 for PdP1, hIP5+hIP8 for PdP2, hIP6+7P for PdP3, and a final grouping that contained 3 regions overlapping PdP4 (hIP6, 7P, 7A). We quantify overlap, with overlap defined as at least one overlapping vertex within each cytoarchitectonic region in a given group and the corresponding anatomical strip centered at each PdP. We first report that each PdP did indeed overlap with hypothesized pairs with high frequency across participants: The strip at PdP1 overlapped the AreahIP7 and AreahIP4 cytoarchitectonic regions (left hemisphere: 92.6% of subjects; right hemisphere: 100%), the strip at PdP2 overlapped the AreahIP5 and AreahIP8 regions (left hemisphere: 72.2% of subjects; right hemisphere: 98.1%), the strip at PdP3 overlapped AreahIP6 and Area7P regions (left hemisphere: 92.6% of subjects; right hemisphere: 90.7%), and the strip at PdP4 overlapped with Area7A (left hemisphere: 85.2% of subjects; right hemisphere: 96.3%), as well as with hIP6 and 7P (**Fig 5A**).

**Figure 5:**
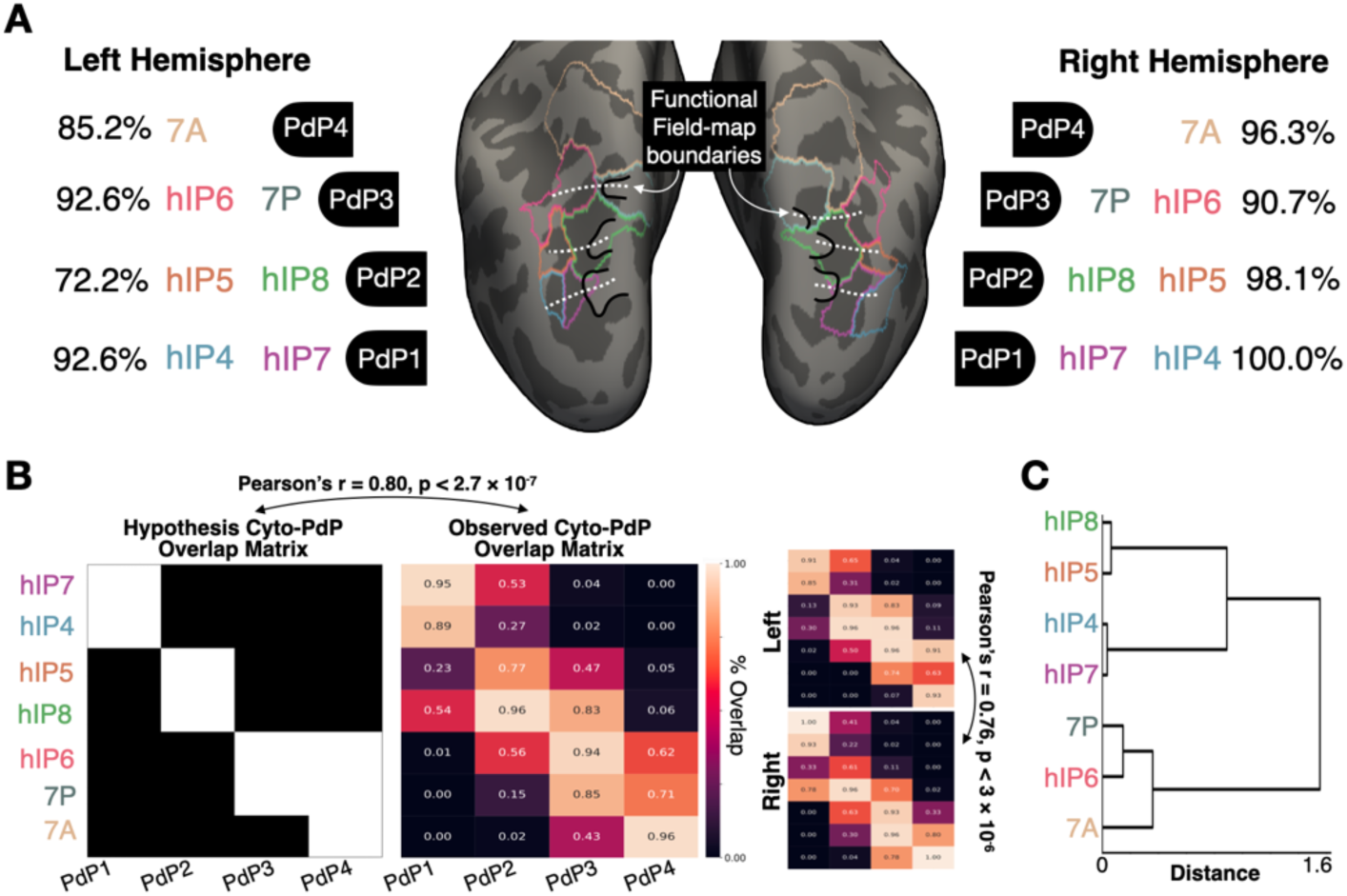
Plis de passage and cytoarchitectonic regions co-localize in human IPS. **(A)** The percentage with which an anatomical strip spanning the IPS-PO defined at each PdP overlaps with single or pairs of cytoarchitectonic regions defined using maximum-probability labels from the Julich Brain Atlas. **(B)** Hypothesis matrix denoting how each PdP (columns) is proposed to overlap with pairs of cytoarchitectonic regions (rows) across individual brains. Cytoarchitectonic labels have been projected into each participant’s native cortical surface where anatomical strips spanning IPS-PO were drawn at each PdP. Matrices on the right depict actual overlap data showing the percentage of hemispheres in which a given PdP and cytoarchitectonic region overlap either bilaterally (larger matrix), or in each hemisphere individually (two smaller matrices on the right). **(C)** Linkage tree denoting clustering and inter-cluster distances from agglomerative clustering performed on bilateral overlap data.

To visualize the preferential overlap, we produce a matrix visualizing the frequency with which a given PdP overlapped one of the cytoarchitectonic regions, averaging performance across hemispheres. Rows and columns were sorted by the order in which each label appears as one ascends the IPS-PO inferiorly to superiorly: PdP1, PdP2, PdP3, PdP4 and AreahIP7, AreahIP4, AreahIP5, AreahIP8, AreahIP6, Area7P, Area7A, respectively, with each cytoarchitectonic medial-lateral pair kept together. We report a significant correlation between the matrix of observed overlap frequency with that of the hypothesis matrix (Pearson’s r = 0.803, p = 2.677 × 10⁻^7^). We find consistency across individual hemispheres as well: The overlap matrix for both hemispheres showed a diagonal pattern, with medial-lateral cytoarchitectonic pairs having the highest overlap with a single PdP (**Fig 5B**). This pattern of overlap between sulcal pleating and cytoarchitecture was significantly correlated across hemispheres (Pearson’s r = 0.758, p = 2.965 × 10⁻^6^). While cytoarchitectonic maps cannot be delineated directly in vivo, the observed overlap patterns suggest intrasulcal pleating of the IPS is consistent with cytoarchitectonic arealization on a broad scale.

While our ordering of cytoarchitectonic regions is qualitative, we can perform agglomerative clustering on this overlap matrix to quantify whether PdP preferentially align with cytoarchitectonic regions and provide quantitative support for our hypothesized medial-lateral pairs. To this end, we performed agglomerative hierarchical clustering on the average left and right hemisphere overlap matrix using average linkage and correlation distance (1 − Pearson’s r), where each individual cytoarchitectonic region was represented by a vector containing its overlap percentages across the four PdP boundaries. We find observer-independent evidence supporting the idea that sulcal pleats overlap specific cytoarchitectonic pairs: AreahIP8-AreahIP5, AreahIP4-AreahIP7, and Area7P-AreahIP6 each clustered together respectively, while Area7A stood alone albeit branching off of the Area7P-AreahIP6 cluster (**Fig 5C**).

Does variability in the pleating and folding of IPS-PO relate to behavior? Sulcal pleating is under strong genetic control given its early gestational emergence (31). In visual cortex, the organization of retinotopic representations is influenced strongly be genetics as well (43), and the cortical sheet continues to fold and change thickness in higher-level visual cortex across development where functional representations mature as children learn to read and recognize (44–47). This raises the possibility that individual variability from PdP-related anatomy may be behaviorally meaningful. We thus ask if features of the cortical sheet within the visual field maps of parietal cortex can capture variance in visual behaviors mediated by the dorsal stream. Given prior work implicating the dorsal visual stream in visuospatial attention (48), we focus on a measure of visuospatial attention bias assessed with a visual line-bisection task (49); **Fig 6A**). We can make two specific hypotheses about how PdP will influence measurements of cortex. First, because gyri tend to be thicker than sulci, we hypothesized that PdP should increase variability in wall thickness, which can be seen along the medial wall of IPS-PO where our PdP exist, but not along the lateral wall (**Fig 3B**). PdP may thus potentially increase standard deviation of cortical thickness. Second, the presence of PdP would likewise introduce convexity into the walls of the sulcus which would otherwise be concave relative to the neighboring gyral ridges. This would reduce the mean curvature (as assessed by FreeSurfer) within IPS-PO. If PdP capture individual variability in sulcal pleating that is behaviorally relevant, then visuospatial behavior should be better predicted by these cortical measures (standard deviation of thickness, mean curvature), rather than by measures that are independent of pleating. For instance, sulcal depth should be insensitive to PdP which protrude perpendicular to IPS-PO depth. Of the adult participants in whom we have anatomically parcellated the dorsal visual stream using PdP, N=27 completed a computerized test of visuospatial bias (**Fig 6A**). From each functionally-defined visual field map (V3AB through IPS-5), we extract the aforementioned cortical properties (Thickness standard deviation, mean curvature, mean sulcal depth) within each participant and average across hemispheres. Cortical features and line-bisection scores were entered into separate GLMs for each anatomical measure, employing a leave-one-out cross-validation approach. Coefficients were estimated from 26 participants and used to estimate the behavior of the held-out participant. As expected, the standard deviation of cortical thickness from the dorsal stream regions, particularly IPS0-IPS3, produced significantly accurate predictions of behavioral visuospatial bias (r = 0.53, p = 0.005) in the expected positive direction. The mean curvature of the dorsal stream regions was also significantly correlated with the true scores (r = -0.55, p = 0.003), showing the expected negative relationship. In contrast, cortical depth produced predictions comparable to chance (r = -0.05, p = 0.800), as expected (**Fig 6C**). These findings suggest that the mesoscale folding features likely influenced by plis de passage relate to dorsal stream-dependent visual behavior.

**Figure 6:**
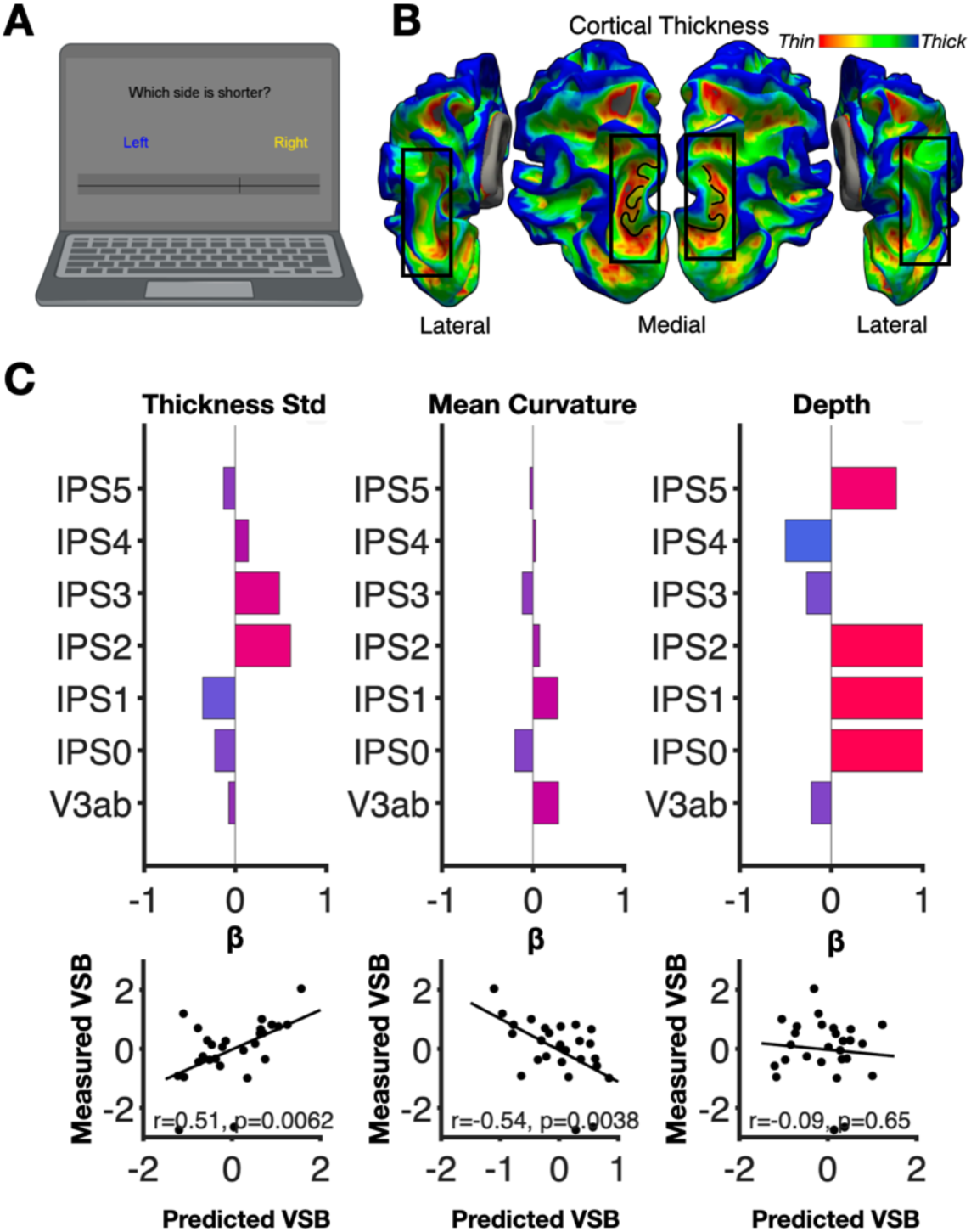
Structural variability introduced by PdP in the cortical sheet of human IPS is correlated with dorsal stream visual behavior. **(A)** Visualization of the computer-based line-bisection test designed to estimate visuospatial bias in degrees of visual angle. **(B)** Cortical surfaces of human brain colored by cortical thickness showing that PdP introduce variability in the thickness of the cortical sheet along the medial wall of IPS-PO. **(C)** Weights and correlation between predicted and measured visuospatial biases across individual adult participants using a leave-one-out cross-validated GLM. Depth, a sulcal metric not impacted by PdP, shows no correlation with behavior. Cortical metrics are derived within each participant’s functionally-defined visual field maps along the IPS-PO.

## Discussion

Our findings extend the early observations by Pierre Gratiolet in 1854 (29), who first reported the intrasulcal pleating phenomenon in macaque parietal cortex, to now show that human parietal cortex contains similar plis de passage that are consistent across individual brains, appear to align with boundaries between functionally-distinct cortex, and may similarly follow the cytoarchitecture of the underlying cortical sheet. Lastly, variability across individuals of cortical folding features introduced by PdP correlate with differences in visual behaviors supported by the dorsal visual stream traversing parietal cortex. These findings are consistent with recent work examining the broader phenomenon of sulcal pleating in humans, which suggested that PdP may account for variability in human sulcal morphology across individuals (32,50). Their conservation across species suggests that PdP may be important markers of brain organization. Moreover, previous work indicates that PdP might also have functional relevance, as they have been linked to the hand motor area (Boling 1999), the temporal voice area (Bodin 2017), and reading behaviors in ventral temporal cortex (34). Despite this potential, PdP remain systematically understudied in humans and absent from modern cortical atlases.

Visual cortex provides a compelling test case to assess the functional and behavioral relevance of this historically-overlooked cortical feature in humans. Increasing evidence suggests that structure–function coupling extends beyond primary sensory areas into higher-order regions, and specifically in the ventral visual stream this coupling has been well established (4–6). In contrast, the dorsal visual stream, a network of regions supporting vision and attention, has lacked an equivalent anatomical framework despite being similarly retinotopic. Here we show that the dorsal stream, which lies buried within a continuous paraoccipital–intraparietal sulcal complex, contains four PdP whose regular and consistent spacing across individuals (**Fig 2**) appears to landmark the approximate boundary separating visual field maps comprising the dorsal stream (**Figs 3-4**). When analyzed in a more observer-independent manner, plis de passage appear to consistently demarcate the interdigitated spacing of foveal and peripheral visual field representations (**Fig 3C**). This further extends the general relationship between structure and function in the human brain to include not just major cortical folds, but pleating features hidden within sulci.

Dorsal stream dysfunction has been implicated in disorders such as ADHD and dyslexia (51–53). One likely hurdle preventing this research is that delineating of the dorsal visual stream currently requires fMRI tasks that demand sustained visual attention during population receptive field (pRF) mapping, which produce visual field maps that require substantial expertise to interpret. These requirements impose steep technical barriers and limit accessibility, particularly in clinical or developmental contexts (Wang et al. 2014; Konen & Kastner 2008; Mackey et al. 2017). As a result, the dorsal stream’s structural–functional organization and developmental trajectory remain largely unknown. The plis de passage characterized here represent a potential neuroanatomical atlas for delineating dorsal stream organization in populations for whom fMRI paradigms are impractical or impossible. The existence of a probabilistic atlas produced from adult and pediatric data (**Fig 4E**) will aid future researchers in identifying the location of PdP within individual brains.

We also observed that each plis de passage of IPS-PO appears to consistently overlap selectively with a pair of cytoarchitectonic regions spanning the medial and lateral banks of the paroccipital sulcus (**Fig 5**). The interpretation of this phenomenon is limited by the fact that cytoarchitectonic estimation was based on data from separate postmortem individuals. We reasoned that if the position of plis de passage within IPS-PO was not conserved across individual brains, then they should have shown high variability in their overlap with cytoarchitectonic regions in an atlas aligned to their native cortical surface. We instead observed preferential overlap that was consistent across individuals and hemispheres (**Fig 5B**). Indeed, prior work has shown that the border between cytoarchitectonic regions of ventral temporal cortex appears in many cases to follow cortical folds, both major sulci such as the occipitotemporal and collateral sulci (54) as well as smaller tertiary sulci such as the mid-fusiform sulcus (55). This prior work is consistent with our observations in that the borders separating medial-lateral pairs as one ascends the IPS-PO occurs in the small sulci between plis de passage, aligning with prior work showing that cytoarchitectonic transitions can occur at sulci in visual cortex. Future work should seek closer homology between cytoarchitecture and cortical folding in the native cortical surface reconstruction of postmortem brains to better understand their relationship.

Lastly, we find that variability in cortical folding characteristics introduced by sulcal pleating captures variance in dorsal stream visual behaviors across individuals. While prior work in ventral visual cortex finds a correspondence between surface are of V1 (56) and V4 (57) with visual behaviors, little work has been done examining the role of sulcal pleating in visual behavior. One study examining the continuity of the occipitotemporal sulcus describes a plis de passage which often bisects the OTS. They find that the presence of a plis de passage within the OTS was associated with higher reading skill, likely contributing to behavioral improvement by providing a cortical surface interface that allows for higher axonal connectivity with the arcuate fasciculus (34). Our work extends this phenomenon to dorsal visual cortex in that increased pleating of the IPS-PO appears to be associated with more balanced visuospatial processing as assessed with a line-bisection task. Our finding that increased pleating of the IPS-PO (higher variance in thickness, more gyral curvature) is associated with more rightward visuospatial scores is in line with prior developmental work showing that young children show a leftward visuospatial bias that becomes more centralized with increasing age (49). Future work analyzing longitudinal data within children should assess if sulcal pleating is established early during infancy or if it is a protracted developmental phenomenon that extends into adolescence and drives the protracted development of reading and visual attention.

In summary, this study identifies a previously unrecognized class of mesoscale cortical folds within the human intraparietal sulcus and demonstrates that they act as conserved anatomical markers for the functional differentiation of the human dorsal visual stream. By revealing a form of structure-function coupling that operates entirely within sulci, these findings extend prevailing assumptions about cortical organization and provide evidence that brain function in humans can be inferred not only across major cortical folds, but from pleating within sulci. A neuroanatomical model linking dorsal stream function to mesoscale structure provides behavioral insights and can be used to parcellate cortex in individuals for whom functional MRI is infeasible.

## Materials and Methods

### Subjects

75 participants were recruited from the broader Mercer Country, New Jersey community. These participants were split into two groups: an observation group (N=33, mean age 30.52 ± 11.71 years, N=13 female) and a replication group consisting of adults and children (Total N=42, ages 7-20, N=30 female; Children N=21, mean age 9.24 ± 1.87, N=15 female; Adults N=21, mean age 18.71 ± 0.78 years, N=15 female). All participants had normal or corrected-to-normal vision and completed structural and functional MRI. A total of 27 subjects from the observation group completed additional behavioral tasks. Written informed consent was obtained in accordance with the Princeton Review Board on human subjects research.

### Structural MRI

MRI data were collected at the Scully Center for the Neuroscience of Mind and Behavior within the Princeton Neuroscience Institute, using a Siemens 3 T Skyra system. Structural MRI was performed for the purposes of reconstructing the cortical surface for data analysis and visualization. Each participant completed T1-weighted (MPRAGE) images: voxel size = 0.8 mm isotropic, TR = 2.4 s, TE = 2.0 ms, flip angle = 8° and T2-weighted images: voxel size = 0.8 mm isotropic, TR = 3.2 s, TE = 565 ms, flip angle = 120°. Both images were used for cortical surface reconstruction in FreeSurfer (v7.4.1), where the T2 image improved gray-white matter segmentation accuracy. Cortical surfaces were reconstructed with FreeSurfer’s recon-all pipeline, generating pial, white matter, and inflated surfaces for each participant. Both individual and averaged analyses were performed. Surfaces were registered to the FreeSurfer average brain for cross-subject comparisons.

### Plis de Passage (PdP) Identification

PdP were manually identified in each hemisphere for every subject using individual native cortical surface reconstructions in FreeSurfer’s FreeView. Major sulcal landmarks were first delineated in order to localise the IPS-PO: anteriorly to posteriorly, the central sulcus precedes the postcentral sulcus, which precedes the IPS, which traverses into its parieto-occipital branch (IPS-PO). Within the IPS-PO, distinct mesoscale pleats were visually identified along the inflated medial wall of the sulcus. Morphologically, when tracing the IPS-PO as it curves around the parieto-occipital fissure (POF), distinct mesoscale pleats emerge from the medial wall and partition the arc of the sulcus into approximately equal thirds (Fig. 2C). This geometric regularity motivated the sequential nomenclature PdP-1, PdP-2, and PdP-3. In addition to these three pleats within the IPS-PO, a fourth, more superficial pli de passage (PdP-4) was identified at the junction separating the IPS-PO branch from the main IPS. All PdP were labeled using a custom annotation protocol implemented in FreeView, and their three-dimensional coordinates were extracted in each subject’s native space.

### PdP Localization

To quantify the spatial consistency of intraparietal PdP, the location of each pleat relative to an anatomical reference was measured. On each participant’s native cortical surface, a point located at the southernmost (posterior and inferior) portion of the IPS-PO was manually identified (Fig. 2E). The shortest path along the cortical surface interconnecting this reference point with a second manually defined point located at the center of each PdP’s gyral protrusion was then derived. From these pairs of points we quantified, in millimeters, the localization of each pleat along the intraparietal sulcus. Statistical differences in PdP location across the four pleats were assessed using a repeated-measures ANOVA (factor: PdP identity, df = 3), conducted separately for each hemisphere.

### Retinotopic Mapping Task

Retinotopic maps were acquired using a modified population receptive field (pRF) paradigm designed to enhance participant engagement and drive higher-level visual responses. During each functional run (300s), participants maintained fixation on a central dot while a bar stimulus (2° visual angle) swept across the visual field along vertical, horizontal, and diagonal axes bidirectionally (8 sweep configurations total). The stimulus covered a circular visual field (radius: 11.5°). Within the moving bar, a stream of colorful cartoon and comic images (>100 unique images) was presented at a rate of 7 Hz. To increase attentional engagement in the dorsal stream, known to elicit attention-dependent responses, the paradigm incorporated a gamified task adapted from Finzi et al. (2021). Participants were instructed to covertly attend to the sweeping bar and detect intermittently appearing target stimuli (animated “bumble bees”) embedded within the broader image stream. The task was framed within a Charlotte’s Web-themed game, in which participants assisted a spider in removing bees from its web. The sweeping bar was presented over a low-contrast radial grid resembling a spider web. Participants responded to detected targets while maintaining central fixation, receiving point-based feedback to encourage sustained attention and task performance. Eye position was continuously monitored using an EyeLink 1000 system (SR Research) to ensure fixation compliance throughout each run.

### fMRI Analysis

Preprocessing followed the Human Connectome Project minimal preprocessing pipeline, including motion correction within-run, slice timing correction (FSL slicetimer), spatial distortion correction (FSL TopUp), co-registration to anatomical image, and surface resampling (FreeSurfer FS-FAST)(58). Functional data were aligned to the acquired T1-weighted images. No spatial or temporal smoothing was applied. Following preprocessing, functional data were analyzed using FreeSurfer’s functional analysis toolbox (FS-FAST). Functional analyses were restricted to the cortical ribbon and conducted in individual native space.

### Population Receptive Field Modeling

Population receptive field (pRF) modeling was performed using a pipeline based on the MrVista implementation (Dumoulin & Wandell, 2008), incorporating a compressive spatial summation (CSS) model (Kay et al., 2013). For each surface vertex, the model estimated a 2D circular Gaussian pRF, yielding the parameters of eccentricity (distance from fixation center in degrees) and polar angle (angular position relative to the upper vertical meridian in degrees). Parameter maps were exported as FreeSurfer overlay files to enable surface visualization and manual delineation of dorsal visual stream regions (V3A–IPS5).

### PdP Alignment with pRF Eccentricity

A strip of cortex was defined that included a given PdP and extended laterally to the opposing wall of the IPS-PO, following the direction of orientation of each PdP in each participant’s native cortical space. Each strip included the three vertices dorsal and inferior from the center of each strip (i.e., six vertices in height; Fig. 3C). From this anatomically defined ROI in each participant, the mean eccentricity value was extracted across pRF fits thresholded at 10% variance explained by the pRF model fit. Peripheral and foveal responses were compared using a one-factor repeated-measures ANOVA with representation (peripheral, foveal) as a within-subject factor. Peripheral values were computed as the mean of the PdP1 (V3AB-IPS0 border) and PdP3 (IPS1-2 border) strips, and foveal values as the mean of the PdP2 (IPS0-1 border) and PdP4 (IPS2-3 border) strips. To examine whether these changes in eccentricity bias emerged simply as a property of the spacing of pleats, the underlying eccentricity map in each participant was permuted (n = 1000) by rotating the map on the cortical surface to preserve spatial autocorrelation in eccentricity tuning.

### PdP Predictive Model of Functional Dorsal Stream

For each participant, one researcher defined the dorsal visual field maps up to IPS-3 to serve as the metric by which anatomical predictions were assessed for accuracy. A separate individual created roughly rectangular anatomical regions of interest (Fig. 4A) to simulate visual field maps. Anatomically defined candidate maps spanned the IPS-PO sulcus, with the medial extent defined by the medial edge of each PdP and the lateral extent defined as the vertex where the sulcal curvature of IPS-PO transitioned to gyral. The anterior–posterior aspect of each anatomically defined map was drawn at the center of each of the four PdP along the IPS-PO. To examine how these anatomical predictions of dorsal visual stream borders compared to the field’s current best model of the dorsal stream, a probabilistic atlas of functionally defined dorsal stream field maps (Wang et al. 2015) was also aligned onto each native cortical surface. This comparison directly evaluated whether anatomy-based predictions could match or exceed probabilistic functional atlases. Because the lower extent of V3AB does not yet have an anatomical definition, the probabilistic version of V3AB (Benson and Winawer 2018) was used in the anatomical model to define the lower boundary of V3AB. For this reason, V3AB was excluded from quantification to maintain statistical independence across datasets.

Atlas prediction performance was evaluated using three metrics. For each ROI, we defined true positive vertices (hits) as the intersection between the functionally-defined label and the model-predicted label, false negatives (misses) as functionally-defined vertices absent from the model prediction, and false alarms as model-predicted vertices absent from the functionally-defined label. False alarms were further subdivided into confusion errors: vertices predicted as the target ROI that belonged to a different dorsal stream ROI, and blank errors: vertices predicted as the target ROI that fell outside all functionally-defined dorsal stream ROIs. Dice coefficient was computed as 2|A∩B| / (|A| + |B|), where A is the functionally-defined, true label and B is the model-predicted label. This metric quantifies spatial overlap between the two label sets, and implicitly penalizes both misses and all false alarms regardless of where they fall on the cortical surface.

Jaccard index was computed as hits / (hits + misses + false alarms), where false alarms include both confusion and blank errors. D-prime (d’) was computed as Z(hit rate) − Z(false alarm rate). D-prime was computed solely within the dorsal stream space to ask a targeted signal detection question: given the set of vertices comprising the dorsal stream ROIs, how well does the atlas discriminate the target ROI from its neighbors? The signal pool was defined as all vertices in the functionally-defined target ROI. The noise pool was defined as all remaining dorsal stream vertices (i.e., vertices belonging to other dorsal stream ROIs). Only confusion false alarms (atlas predictions landing on vertices truly belonging to another dorsal stream ROI) contributed to the false alarm rate, as blank errors fall outside the defined noise pool. Hit and false alarm rates were adjusted using the log-linear correction to avoid infinite z-scores.

Boundary distances were computed in each subject’s native surface space by extracting the 3D coordinates of vertices belonging to each boundary label (true functional, predicted anatomical, and predicted probabilistic) from the FreeSurfer white matter surface. Centroid distance was calculated between each predicted boundary and the corresponding true boundary, defined as the Euclidean distance between the mean coordinates of each boundary’s vertex set. To compare boundary distances between the anatomical and probabilistic predictions, we used a repeated measures ANOVA with AtlasType (anatomical, probabilistic), Boundary (V3ab/IPS0, IPS0/IPS1, IPS1/IPS2, IPS2/IPS3), and Hemisphere (left, right) as within-subject factors. Analyses were conducted in MATLAB using the fitrm and ranova functions.

### PdP and Cytoarchitecture Correspondence

Subject-specific anatomical strips centered at each PdP spanning the IPS-PO were defined, as described in the PdP Alignment with pRF Eccentricity methods section. Cytoarchitectonic labels were obtained from the Julich Brain Atlas and projected into each participant’s native cortical surface space. Seven regions spanning the IPS-PO complex were analyzed: hIP4, hIP5, hIP6, hIP7, hIP8, Area 7P, and Area 7A. For each participant and hemisphere, we assessed whether each PdP strip overlapped a given cytoarchitectonic region, defining overlap as the presence of at least one shared surface vertex. Based on the medial-lateral organization of the cytoarchitecture underlying the IPS-PO complex, we grouped regions into hypothesized pairs corresponding to each PdP: hIP4+hIP7 (PdP1), hIP5+hIP8 (PdP2), hIP6+Area 7P (PdP3), and a final grouping consisting of hIP6, Area 7P, and Area 7A (PdP4). For each PdP-cytoarchiture pairing, we computed the proportion of participants exhibiting overlap. To test for preferential correspondence, we constructed an observed overlap matrix reflecting the frequency with which each PdP overlapped each cytoarchitectonic region, averaged across hemispheres. Rows and columns were ordered according to the inferior-superior progression along the IPS-PO (row order: hIP7, hIP4, hIP5, hIP8, hIP6, Area 7P, Area 7A; column order: PdP1,PdP2, PdP3, PdP4). The observed matrix was compared with the hypothesis matrix using Pearson correlation (Fig.5B). Hemispheric consistency was evaluated by computing overlap matrices separately for left and right hemispheres and correlating the resulting matrices using Pearson correlation (Fig. 5C).

To quantify whether PdP-defined cortical boundaries preferentially align with specific cytoarchitectonic regions, we performed agglomerative hierarchical clustering separately for the left and right hemispheres. For each hemisphere, we constructed a cytoarchitectonic × PdP overlap matrix in which each cytoarchitectonic region was represented by a vector of overlap percentages across the four PdP strips. Prior to clustering, cytoarchitectonic regions with no overlap across all PdP were excluded. Remaining missing values were imputed using the column-wise mean to enable computation of pairwise distances. Pairwise dissimilarity between cytoarchitectonic regions was computed using correlation distance (1 − Pearson correlation), and agglomerative clustering was performed using average linkage. Clustering was applied independently to the left and right hemisphere matrices.

### GLM Prediction of Visuospatial Bias from Dorsal Stream Cortical Properties

Of the adult participants in whom the dorsal visual stream was anatomically parcellated using PdP, N = 27 completed a computerized line-bisection task (Fig 6A). From each functionally defined visual field map (V3AB through IPS-5), the standard deviation of cortical thickness and mean curvature were extracted using FreeSurfer’s mris_anatomical_stats function, and sulcal depth was extracted using FreeSurfer’s recon-all .sulc file, within each participant and averaged across hemispheres. Cortical features and line-bisection scores were entered into separate GLMs for each anatomical measure, employing a leave-one-out cross-validation approach. Coefficients were estimated from 26 participants and used to predict the behavior of the held-out participant.

## Code and Data Availability

Code and data related to figure reproduction can be found at https://github.com/anna-lyn-williams/PlisDePassage

## Acknowledgements

This work was supported by funding from the National Institute of Health’s National Eye Institute (nei.nih.gov) R01EY036881 to JG, as well as funding from the Whitehall Foundation award 2023-08-052 (whitehall.org). This work was also supported in part by a National Science Foundation (nsf.gov) GRFP award to ALW, and Gilliam Fellowship funding from the Howard Hughes Medical Institute (hhmi.org) to PMH. Funders played no role in the design of the experiment, collection or analysis of the data, decision to publish, or preparation of the manuscript.

## Notes

### Competing Interest Statement

The authors have declared no competing interest.

